## Supplemental Figures for "The type 3 secretion system requires actin polymerization to open translocon pores"

### Supplemental Figure Legends

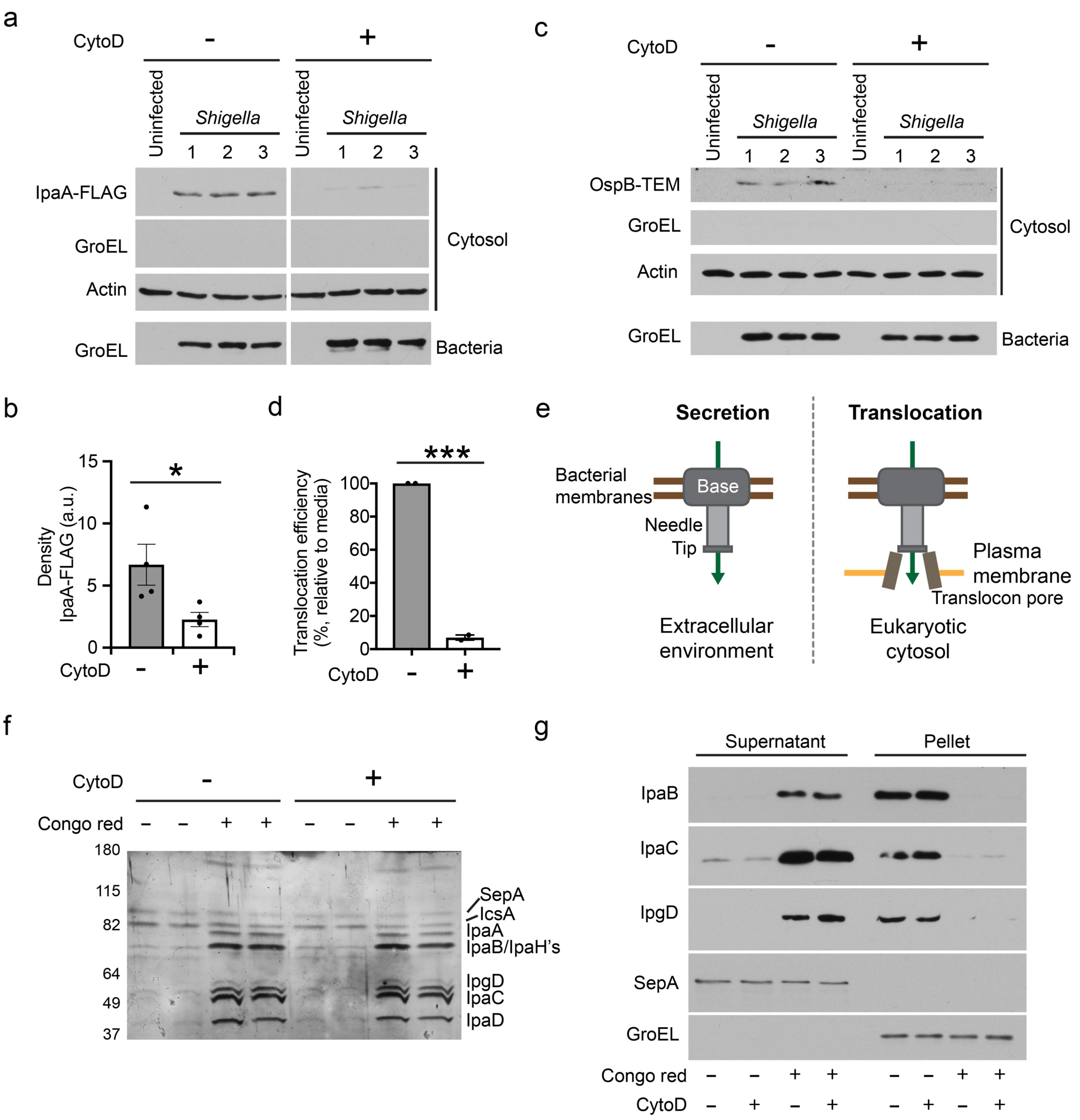


**Figure S1. T3SS effector translocation requires actin polymerization.**

(a-d) *S. flexneri* translocation of the FLAG-tagged type 3 effector IpaA and the TEM β-lactamase-tagged type 3 effector OspB into HeLa cells requires actin polymerization. (a and c) Representative western blots of cytosolic FLAG-tagged IpaA (a) or TEM-tagged OspB (c) in *S. flexneri* infected HeLa cells. GroEL, bacterial cytosolic protein; actin, eukaryotic cytosolic protein. (b and d) Quantification of cytosolic effectors FLAG-tagged IpaA (b) and TEM-tagged OspB (d) from experiments depicted in panels a and c, respectively. *, $p<0.05$; ***, $p<0.001$; Student's t-test. (e) Schematic diagram showing differences between induced type 3 mediated secretion from bacteria in liquid media, which results in bacterial effectors in the extracellular medium, and plasma membrane contact-induced type 3 mediated translocation, which results in bacterial effector protein translocation into the host cytosol. (f-g) Effect of CytoD on type 3 secretion of effectors following induction with Congo red. (f) Bacterial supernatant proteins detected in silver-stained gel. Representative of four experiments. (g) Western blots of bacterial supernatants and pellets, representative of three independent experiments. IpaA, IpaB, IpaC, IpaD, IpaH's, and IpgD, type 3 secreted proteins. GroEL, bacterial cytoplasmic protein; SepA and IcsA, type 5 secreted proteins, whose secretion occurs independent of both the T3SS and Congo red.

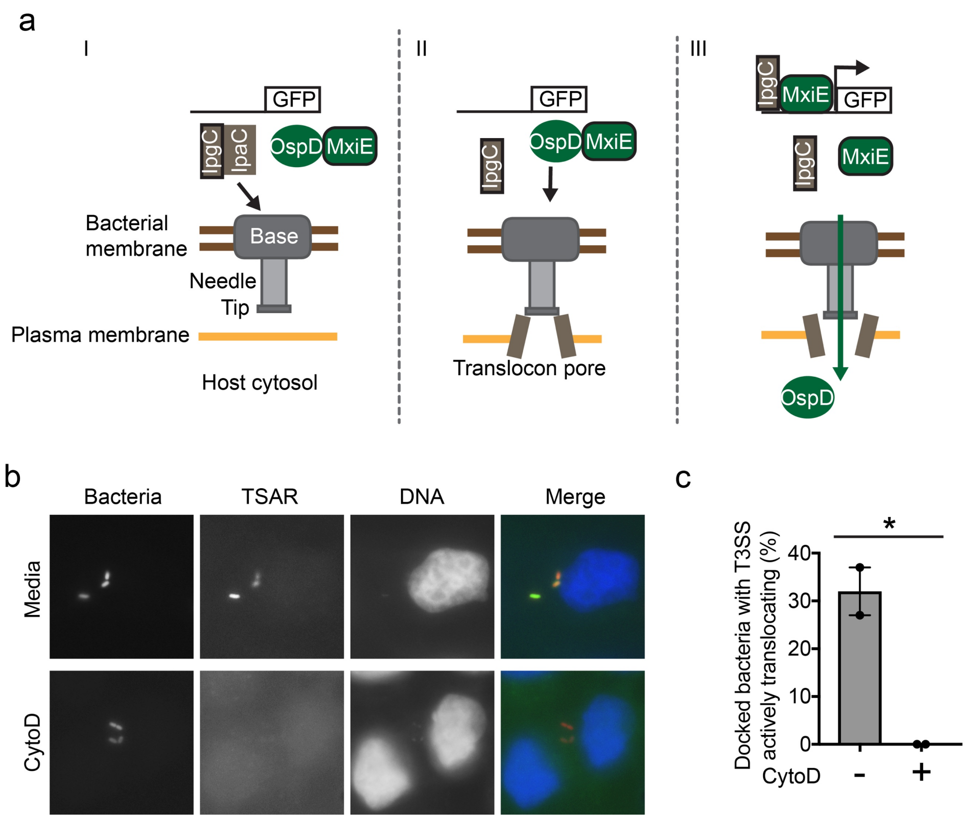


**Figure S2. Type 3ecretion activity, as measured by TSAR, requires actin polymerization.**

(a) Schematic depiction of OspD-dependent production of GFP by the TSAR reporter. Transient contact with host plasma membrane activates secretion of the translocon pore proteins IpaC and IpaB (IpaB not shown) (I). The secretion of IpaC and IpaB liberates their cognate chaperone, IpgC, and secreted IpaB and IpaC form the translocon pore in the plasma membrane, onto which the bacterium docks (II). OspD translocation liberates its chaperone, MxiE, IpgC binds to and activates MxiE, and IpgC-MxiE function as a transcriptional activator, inducing the *mxiE* promoter upstream of *gfp* (III). (b) Infection of HeLa cells with *S. flexneri* carrying TSAR. Representative fluorescent images. Blue, DNA (Hoechst); red, mCherry (constitutively produced); green, GFP (transcriptionally activated by the secretion of OspD). (c) Percentage of docked bacteria with active secretion in experiments represented in panel b. Data points represent independent experiments. *, $p<0.05$; Student's t-test.

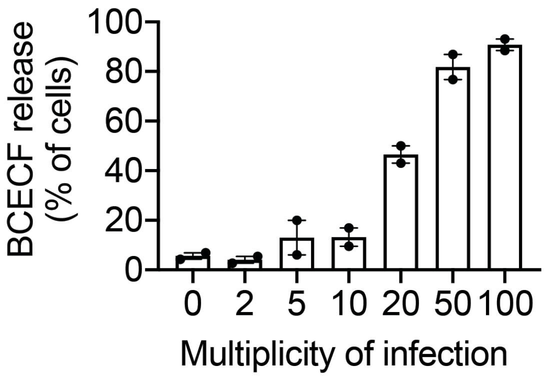


**Figure S3. Formation of intermediary pores in the plasma membrane does not require actin polymerization.**

BCECF dye released from HeLa cells infected with *E. coli* pSfT3SS as a function of multiplicity of infection. Data are mean ± SEM of two independent experiments; data points are independent experiments.

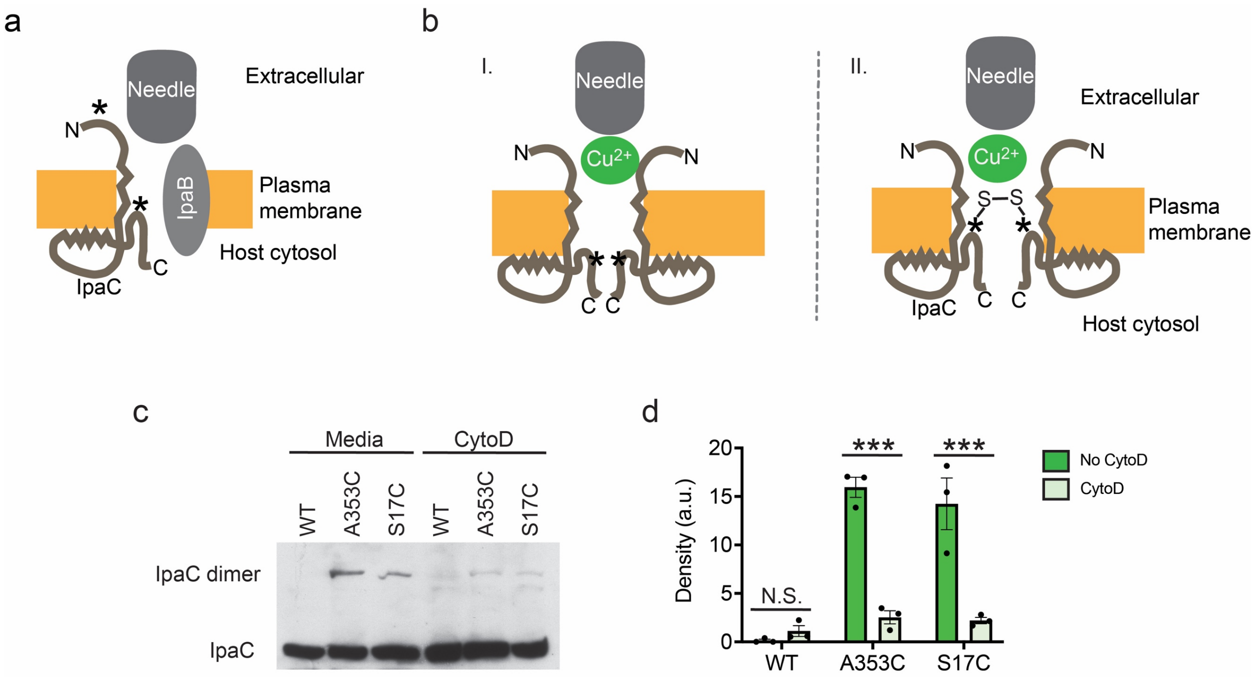


**Figure S4. Copper oxidant mediated crosslinking of IpaC requires actin polymerization.**

(a) Schematic depiction of the position of S17C and A353C in IpaC indicated by asterisk. (b) Schematic depiction showing that disulfide bonds do not form between adjacent IpaC monomers when A353C is in the cytosol (I), but can form in an intermediary pore complex when an A353C-containing loop of IpaC extends into the lumen of the pore (II). (c-d) Effect of cytoD on the ability of the oxidant copper to induce crosslinking between IpaC monomers at S17C and A353C. (c) Representative western blot. (d) Quantification of crosslinked dimer band density in panel d. Data are the mean ± SEM of three independent experiments. Data points are independent experiments. ***, $p<0.001$ by two-way ANOVA with Sidak *post hoc* test.

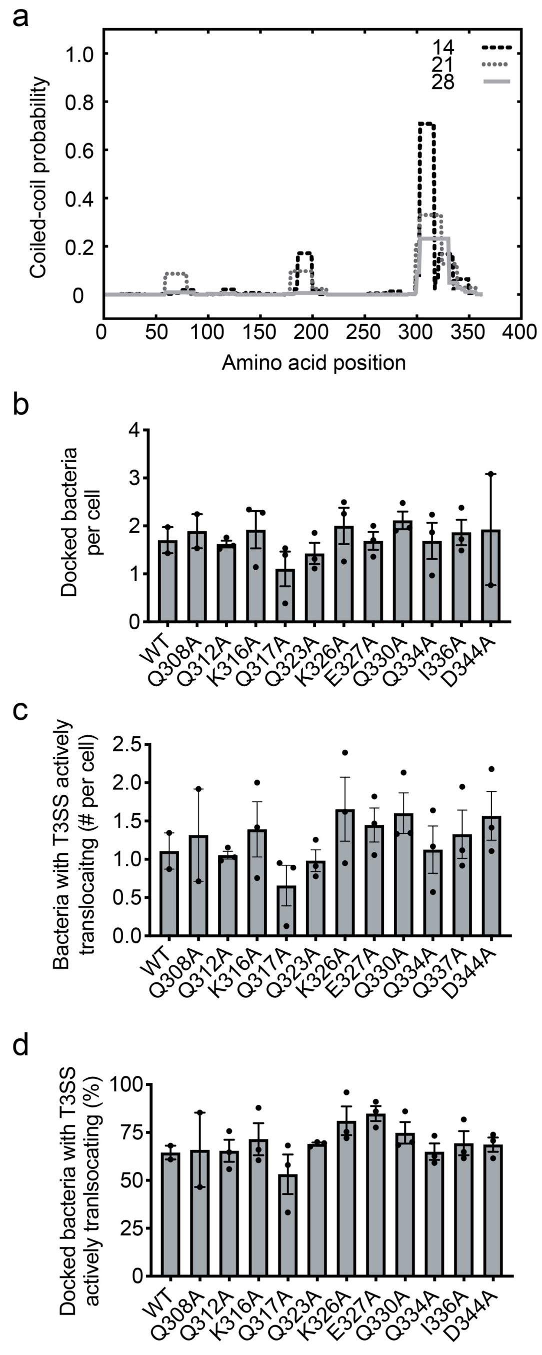


**Figure S5. Scanning alanine mutagenesis of charged/polar residues in the coiled-coil domain did not identify individual residues critical for docking or effector translocation.**

(a) Prediction of coiled-coil domains in IpaC by COILS [46]; 14, 21, and 28 indicate the number of amino acids in each coil. (b-d) Docking and translocation into MEFs by *S. flexneri* strains producing indicated IpaC alanine mutant. (b) Docked bacteria per cell at 50 minutes of infection. (c) Number of bacteria with active secretion per cell. (d) Percentage of docked bacteria with active secretion. (b-d) Data are mean ± SEM from two to three independent experiments; data points represent individual experiments.

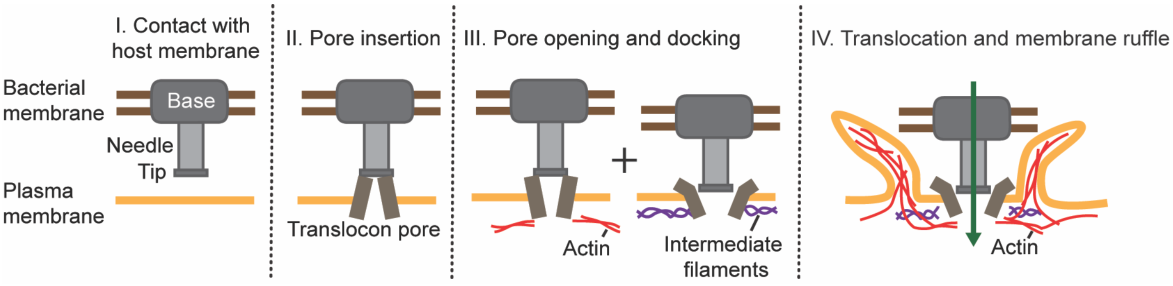


**Figure S6. Model.**

Contact of the T3SS with the host plasma membrane (I) induces the T3SS to deliver the translocon pore proteins into the plasma membrane (II). Actin polymerization opens the pore, and the interaction of IpaC with intermediate filaments promotes bacterial docking onto the pore complex (III). Effectors are secreted through the T3SS, and together with IpaC, trigger membrane ruffle formation (IV) and consequent bacterial uptake.
